# Prediction and evolution of the molecular fitness of SARS-CoV-2 variants: Introducing SpikePro

**DOI:** 10.1101/2021.04.11.439322

**Authors:** Fabrizio Pucci, Marianne Rooman

**Affiliations:** Computational Biology and Bioinformatics, Université Libre de Bruxelles, Brussels, Belgium; Interuniversity Institute of Bioinformatics in Brussels, Brussels, Belgium

**Keywords:** Viral Fitness, SARS-CoV-2, CoViD-19, Spike protein, Variants, Mutagenesis, Folding Free Energy, Binding Free Energy

## Abstract

The understanding of the molecular mechanisms driving the fitness of the SARS-CoV-2 virus and its mutational evolution is still a critical issue. We built a simplified computational model, called SpikePro, to predict the SARS-CoV-2 fitness from the amino acid sequence and structure of the spike protein. It contains three contributions: the viral transmissibility predicted from the stability of the spike protein, the infectivity computed in terms of the affinity of the spike protein for the ACE2 receptor, and the ability of the virus to escape from the human immune response based on the binding affinity of the spike protein for a set of neutralizing antibodies. Our model reproduces well the available experimental, epidemiological and clinical data on the impact of variants on the biophysical characteristics of the virus. For example, it is able to identify circulating viral strains that, by increasing their fitness, recently became dominant at the population level. SpikePro is a useful instrument for the genomic surveillance of the SARS-CoV-2 virus, since it predicts in a fast and accurate way the emergence of new viral strains and their dangerousness. It is freely available in the GitHub repository github.com/3BioCompBio/SpikeProSARS-CoV-2.

## 1. Introduction

Despite mitigation measures put in place around the world to slow down the fast spreading of the SARS-CoV-2 virus, the CoViD-19 viral pandemic continues to have global devastating effects, with more than 130,000,000 people infected and almost 3,000,000 deaths [1]. Lots of efforts and resources have been devoted in the last year to develop vaccines and new therapeutics in response to the SARS-CoV-2 infection [2,3]. Several vaccines such as mRNA-1273 [4], BNT162b2 [5], AZD1222 [6], Sputnik V [7], Ad26 [8] and NVX-CoV2373 [9] have proven to be safe and efficacious against the viral agent and have recently been approved by the regulatory agencies for emergency use. Thanks to these developments, large-scale vaccine administration is now ongoing throughout the world.

Moreover, while the pathogenic mechanisms of the viral infection are still unclear, effective therapeutic agents have been developed. For example, neutralizing antibodies (nAbs) targeting the viral spike protein or human convalescent plasma have been employed in clinical practice by passively transferring them to patients [10–13]. This therapy generally leads to an improvement of the disease conditions and to a reduction of viral load.

The increase in viral immunity at the population level due to infection, vaccination or passive immunization via nAbs clearly results in a stronger selection pressure on the SARS-CoV-2 virus [14,15]. This causes the emergence of new variants of the virus which are able to escape from the immune response. Lots of computational and experimental studies are currently focusing on the understanding of these escape mechanisms in the SARS-CoV-2 viral infection [16–20] and on setting up SARS-CoV-2 immune surveillance of the world’s population to track and eventually limit the spreading of potentially escaping variants [21–26].

However, the prediction of how SARS-CoV-2 evolves under this selective pressure is far from obvious. Indeed, even though SARS-CoV-2 has a moderate mutation rate compared to other RNA viruses due to its more accurate replication [27], tracking viral dynamics in the huge space of possible variant combinations (including also deletions and insertions) under the influence of human immunity makes predictions highly challenging. Extensive large-scale monitoring of SARS-CoV-2 evolution and host immunity will help to better understand these issues [27].

In this paper, we performed an extensive computational analysis of the mutational mechanisms that lead to the emergence of SARS-CoV-2 strains with increased fitness, with the aim to better understand the molecular mechanisms that drive viral adaptation and escape from the human immune system. We performed *in silico* mutagenesis experiments and predicted the impacts of mutations in the spike protein on its stability and on its affinity for nAbs and for the angiotensin-converting enzyme 2 (ACE2), known to be the SARS-CoV-2 entry point into the cell. We validated these predictions on viral variants for which experimental, epidemiological or clinical data has been obtained, and especially on the variants that are emerging and rapidly spreading to become prevalent genotypes. Our predictions are of utmost importance to help monitoring the future evolutionary dynamics of SARS-CoV-2 and to identify the emergent strains whose spread will have to be limited via either the design of new vaccines or new mitigation measures.

## 2. Methods

### 2.1. Spike protein structures

The spike protein or S-protein of the SARS-CoV-2 virus (Uniprot code P0DTC2) is a homotrimeric glycoprotein attached to the viral membrane. It can adopt two forms, a closed and an open form. The transition between these forms increases the solvent exposure of the protein’s receptor-binding domain (RBD), which encompasses residues 333-526 and mediates the fusion of the membranes of the virus and its host.

The 3-dimensional (3D) structures of the two forms have been experimentally resolved by cryo-electron microscopy (cryo-EM) and are deposited in the Protein DataBank (PDB) [28]. The closed form, with PDB code 6VXX, has a resolution of 2.80 Å [29], and the open form, 6VYB, has a resolution of 3.20 Å. These structures have thus quite a low resolution and do not contain all the residues of the spike protein. To get structures of the closed and open forms without missing residues, we modelled the complete amino acid sequence using the PDB structures 6VXX and 6YVB as templates and the homology modelling webserver SWISS-MODEL [30].

More accurate structures, resolved by X-ray crystallography, are available for the RBD of the spike protein. We used the PDB structure 6M0J [31] for this region, which contains the RBD bound to ACE2, with a resolution of 2.45 Å.

Furthermore, we set up a dataset of spike protein/nAb complexes taken from [32], referred to as 𝒟 ^nAb^. We used the following selection criteria:

- Human monoclonal nAbs generated in response to SARS-CoV-2 infection;
- nAbs targeting the spike protein;
- nAbs/spike protein complexes available in the PDB, with X-ray structure of resolution ≤ 3.2 Å.

𝒟 ^nAb^ contains 31 structures of nAbs/spike protein complexes, listed in the GitHub repository github.com/3BioCompBio/Sp CoV-2. These nAbs exclusively target the RBD of the spike protein, and are assumed to mimic the diversity of the human immune B-cell repertoire.

### 2.2. Spike protein stability

To compute the change in folding free energy upon point mutations in the spike protein, we used the PoPMuSiC algorithm [33], which is based on the 3D structure of the target protein and a combination of statistical mean-force potentials. We applied it to the modelled structures of the open and closed forms of the spike protein, and to the experimental structure of the RBD domain. The final value of the change in folding free energy caused by a mutation *i*, 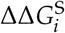, was defined as follows: for mutations of residues in the RBD, we considered the predictions based on the 6M0J structure of the RBD; for mutations of other residues, we averaged the predicted energy values obtained from the two models obtained from the low-resolution structures 6VYB and 6VXX.

### 2.3. Spike protein/ACE2 binding affinity

For the changes in binding affinity upon single-site mutations, we used the BeAtMuSiC predictor [34], which is a linear combination of free energy values predicted by PoPMuSiC on the protein complex and on the separate partners. We applied BeAtMuSiC to predict the effect of variants in the viral spike protein on its binding affinity for the ACE2 receptor of the host, which allows entry of SARS-CoV-2 virus into cells. For this purpose, we considered the X-ray structure 6M0J of the RBD/ACE2 receptor complex [31] as input, and computed the change in binding free energy 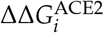 of the RBD/ACE2 complex upon mutations *i* in the RBD. Mutations in the spike protein but outside of RBD were assumed to have no effect on ACE2 binding.

### 2.4. Spike protein/nAb binding affinity

The changes in binding affinity between the spike protein and the 31 nAbs from the 𝒟^nAb^ set caused by point mutations in the spike protein were also estimated using BeAtMuSiC [34]. We computed the effect of each mutation *i* on the binding affinity 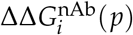 of each nAb/spike protein complex p, and computed their mean value over the 31 complexes from 𝒟^nAb^:

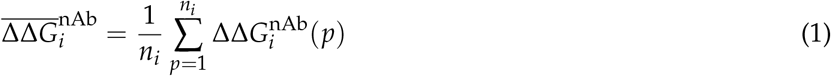

where *n*_*i*_ is the number of structures that include the mutation *i*. Indeed, the structures of the nAb/spike protein complexes do not cover exactly the same region of the spike protein.

### 2.5. SARS-CoV-2 fitness

Viral fitness is related to how efficiently the virus produces infectious progeny [35]. It is a fairly complex function of different characteristics among which the transmissibility of the virus, its infectivity and its ability to escape from the host’s immune response. We estimated the fitness Φ_*i*_ of a variant *i* of the SARS-CoV-2 virus on the basis of a simplified model which only takes into account the spike protein. More precisely, we defined it as a product of three fitness contributions:

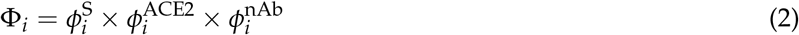

where *ϕ*^S^, *ϕ*^ACE2^ and *ϕ*^nAb^ represent the relative propensities of the mutant virus to be transmitted, to infect the host, and to escape the host’s immune system. These propensities are assumed to be higher for spike protein variants that are stabler [36] (ΔΔ*G*^S^ < 0), that have greater binding affinity for the ACE2 receptor [37] (ΔΔ*G*^ACE2^ < 0), and that have lower binding affinity for nAbs 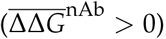, respectively. We thus defined the fitness contributions 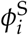 and 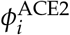 of a mutation *i* to be a positive decreasing function of 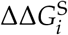 and 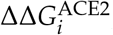, respectively, and 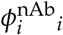a positive increasing function of 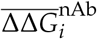. More precisely:

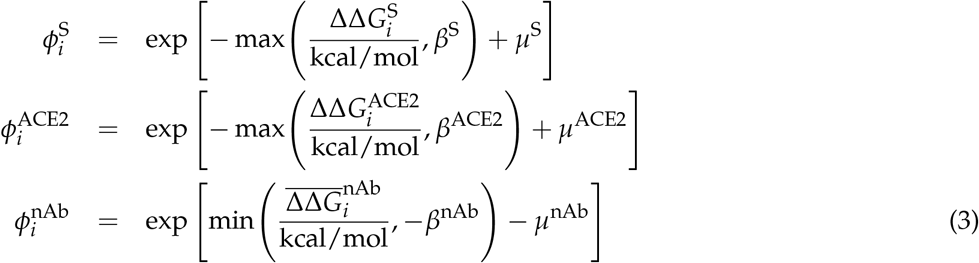

where *µ*^S^, *µ*^ACE2^, *µ*^nAb^, *β*^S^, *β*^ACE2^ and *β*^nAb^ are parameters. The choice of the *ϕ*-functions and parameters is justified as follows:

- Mutations *i* that strongly destabilize the spike protein 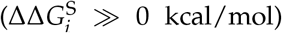 or its binding to ACE2 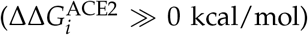, or that stabilize its binding with nAbs 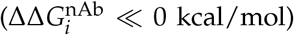 have a fitness close to zero.
- Mutations that stabilize the spike protein 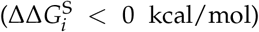 or its binding to ACE2 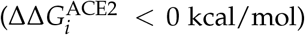, or that destabilize binding to nAbs 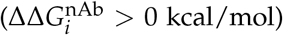 have an evolutionary advantage and a fitness higher than one.
- To avoid excessively high fitness values, we cut the exponential growth of the *ϕ*-functions for ΔΔ*G*_*i*_ = *β*, with *β* = *β*^S^ = *β*^ACE^ = *β*^nAb^ chosen to be −1, similarly to what has been proposed in [38].
- The folding free energy changes predicted by PoPMuSiC have been shown to be biased towards destabilizing mutations [39,40]. To correct for this effect, the *µ*^S^ parameter has been chosen to be equal to 0.5. The changes in binding free energy predicted by BeAtMuSiC have an analogous bias, as they are constructed from PoPMuSiC scores. Following the BeAtMuSiC construction detailed in [34], a bias in the PoPMuSiC energy value of 0.5 results in a bias in the BeatMuSiC energy value of 0.19. We thus fixed *µ*^S^ = 0.50 and *µ*^ACE^ = *µ*^nAb^ = 0.19.
- We set by definition the fitness value of the wild-type equal to one: 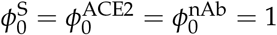.

The global viral fitness, which takes into account multiple mutations in the spike protein, is defined as the product of the fitness values of all point mutations *i* as:

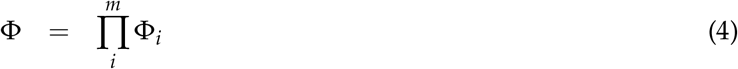

where *m* correspond to the total number of mutations in the spike protein relative to the wild-type strain. Note that, in doing so, we considered the mutations as independent and discard possible epistatic effects.

## 3. Results

### 3.1. Computational pipeline

In its viral evolution, SARS-CoV-2 and our immune system are constantly engaged in what is known as a cat-and-mouse game, where SARS-CoV-2 attempts to increase its fitness by increasing its transmissibility, infectivity and/or to escape from the human immune response. To quantitatively describe the viral fitness landscape, we developed a simplified model in which we focused on the spike protein. This protein, which protrudes from the virus surface, is a crucial component of the infection, as its binding to the ACE2 receptor of the host mediates the virus entry into the cells. The binding affinity of the spike protein for ACE2 has thus been related to SARS-CoV-2 infectivity [37]. The stability properties of the spike protein itself are another key element in the viral infection which has been related to the viral transmissibility [36].

Moreover, the spike protein is a major inducer of the host’s immune response [18,26]. We mimicked the effect of the immune system on the SARS-COV-2 virus through a set of 31 nAb/spike protein complexes contained in the dataset 𝒟 ^nAb^ (see Section 2.1). We observed that these nAbs target exclusively the RBD of the spike protein and that the epitopes cover almost the entire RBD surface, as shown in Fig. 1. A recent investigation suggests that RBD-binding antibodies are the major contributors of the neutralizing activity in convalescent human plasma [18,26]. This justifies our approximation of considering the nAbs of the set 𝒟 ^nAb^ as representative of the immune response.

**Figure 1.**
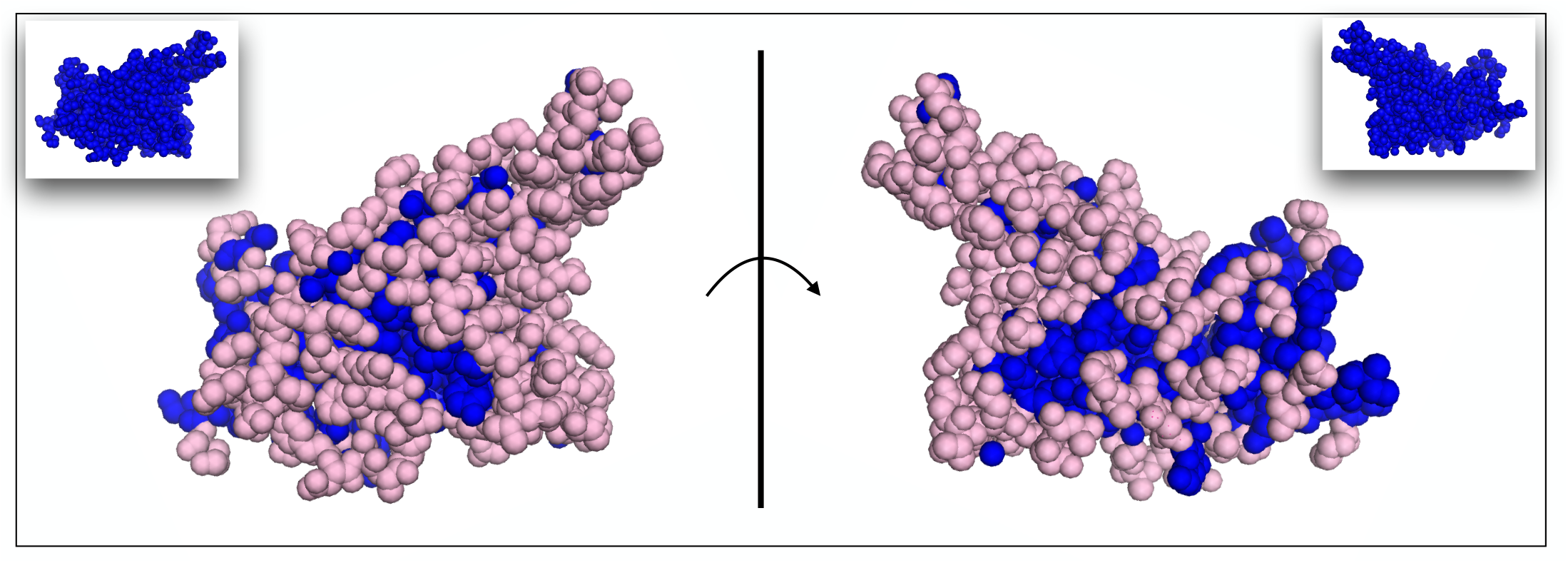
Receptor binding domain of the SARS-COV-2 spike protein. The ensemble of epitope residues targeted by at least one of the nAbs of the set 𝒟 ^nAb^ are in light pink spheres, and the other, non-epitope, residues are in blue spheres. The two views are related by a 180^*o*^ rotation with respect to the plane representing the ACE2 binding interface.

To estimate the global viral fitness F of spike protein variants in terms of transmissibility, infectivity and escape from the host’s immune response, we computed it, using physics-based approaches, as a product of three fitness contributions, related to the change in stability of the spike protein upon amino acid substitution (*ϕ*^S^), and to its change in binding affinity for ACE2 (*ϕ*^ACE2^) and for neutralizing antibodies (*ϕ*^nAb^), respectively, as defined in Eqs (1)-(3). The effect on fitness of multiple mutations are considered as independent and thus simply multiplied (Eq. 4). The fitness contributions are in turn expressed in terms of the change upon mutation of the folding free energy of the spike protein (ΔΔ*G*^S^) and of its binding affinity for ACE2 (ΔΔ*G*^ACE2^) and for nAbs (ΔΔ*G*^nAb^), using the PoPMuSiC [33] and BeAtMuSiC [34] algorithms (see Sections 2.2-2.4).

In order to identify mutations in the spike protein that increase or decrease the SARS-CoV-2 transmissibility or infectivity, or that facilitate or block the escape from the protective immunity elicited by the infection, we constructed a computational pipeline of three steps, schematically represented in Fig. 2, in which we estimated ΔΔ*G*^S^and *ϕ*^S^, ΔΔ*G*^ACE2^ and *ϕ*^ACE2^, and ΔΔ*G*^nAb^ and *ϕ*^nAb^. Using this pipeline, we performed large-scale computational mutagenesis experiments, in which we introduced basically all mutations in the spike protein and predicted their effect on viral fitness. In what follows, we confronted these predictions with a large series of available experimental, epidemiological and clinical data on the SARS-CoV-2 infection and evolution.

**Figure 2.**
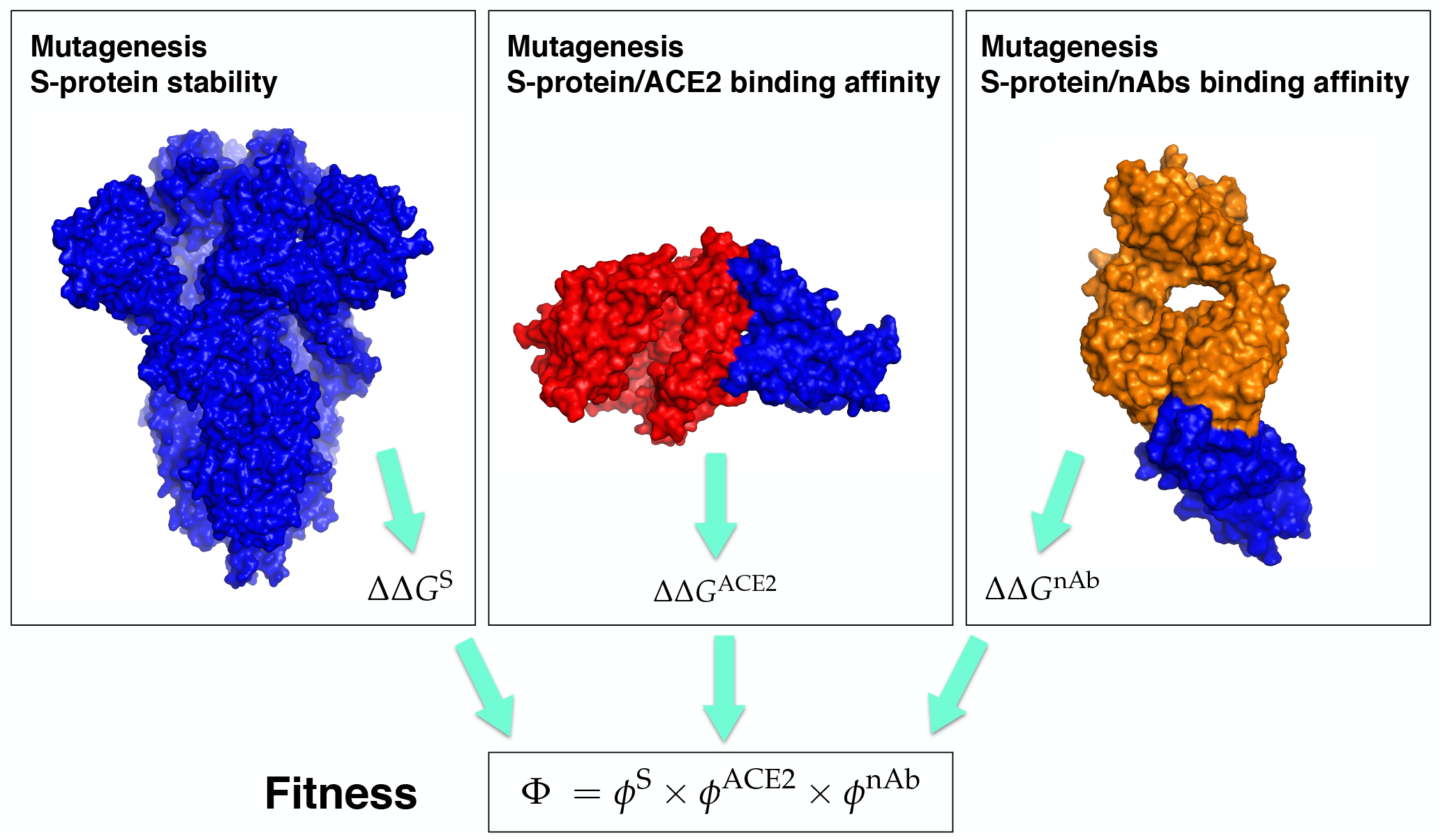
Schematic representation of the three steps of our computational pipeline: *in silico* mutagenesis experiments to compute the change in stability of the spike protein, and its change in binding affinity for ACE2 and for the 31 nAbs from 𝒟^nAb^. The spike protein is in blue, ACE2 in red and the antigen-binding fragment of a nAb in orange. The structures used for the pictures on the left, center and right have the PDB codes 6VXX, 6M0J and 7B3O, respectively.

Our prediction pipeline, called SpikePro, is freely available as an easy-to-use c++ program, which needs a variant spike protein sequence in fasta format as input. It outputs the sequence alignment with the reference spike protein (Uniprot code P0DTC2), the list of all point mutations introduced and the predicted overall viral fitness F. It can be downloaded from github.com/3BioCompBio/SpikeProSARS-CoV-2.

### 3.2. Spike protein stability and SARS-Cov-2 transmissibility

We performed a large *in silico* mutagenesis experiment to study the influence of mutations on spike protein stability and thus on viral transmissibility [36]. Using PoPMuSiC [33], we computed the change in folding free energy 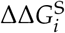 of all possible single-site mutations *i* in the spike protein, and the corresponding fitness contribution 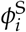 *i* defined in Eq. (3).

As a first check of our method, we analyzed the relation between the predicted 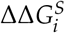 values for all point mutations in the RBD domain and the measured effects of these variants on the spike protein expression [41]. These measurements were done using a yeast surface display platform, in which protein expression was quantitatively determined at large scale via flow cytometry. Even though protein expression and stability are only partially correlated, we found a good Pearson correlation coefficient of −0.51 between the measured expression and the predicted 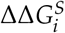 values, which can be considered as the first validation of our approach.

To analyze the relation between stability predictions and epidemiological data, we compared the computed spike protein stability changes 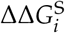 with the observed mutation rate *R*_*i*_. We estimated *R*_*i*_ as the number of occurrences of each point mutation *i* in the set of about 7.8 × 10^5^ SARS-CoV-2 spike protein sequences collected in the GISAID database [42], divided by the number of residues in the spike protein. We analyzed *R*_*i*_ as a function of the predicted 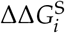 values for all possible mutations *i* in the whole spike protein. As seen in Fig. 3.a, the majority of mutations that became dominant during the evolutionary trajectory show a slight increase of the spike protein stability, with 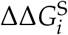 between −1 and 0 kcal/mol. A smaller number of dominant variants have their stability slightly decreased with 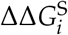 between 0 and 1 kcal/mol. Outside of this free energy interval, the rate *R*_*i*_ is almost vanishing.

**Figure 3.**
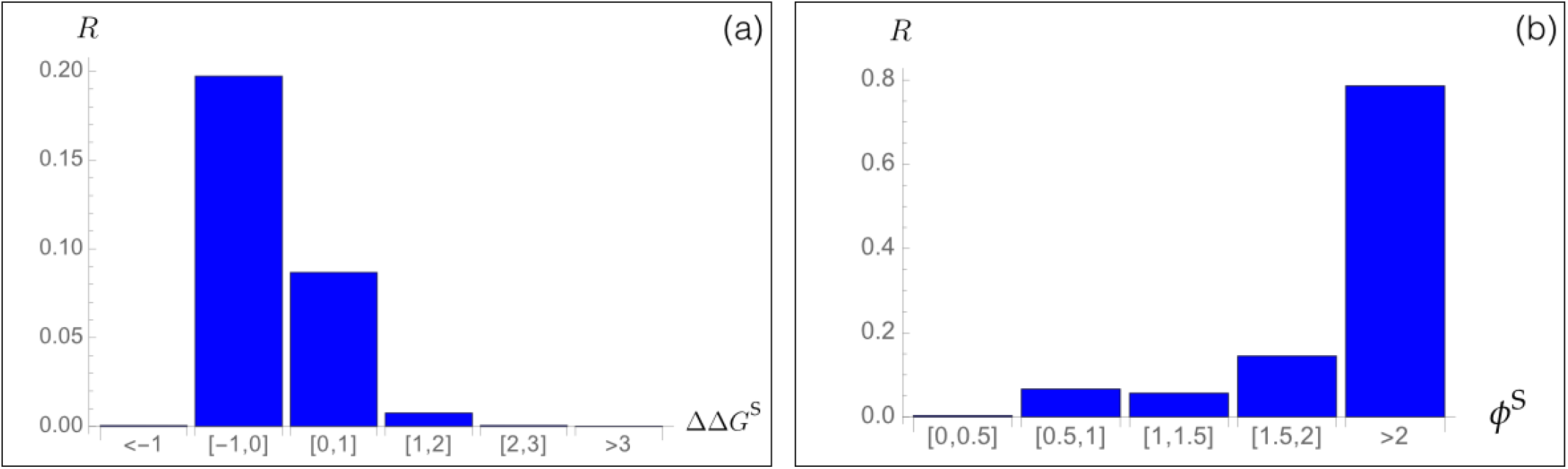
Mutation rate *Ri* of SARS-CoV-2 spike protein variants *i* observed in the GISAID database [42] as a function of (a) the predicted change in folding free energy 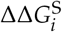 (in kcal/mol) and (b) the predicted fitness contribution 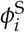.

Moreover, we found a very good agreement between the predicted fitness 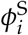 and the *R*_*i*_ rate, as seen in Fig.3.b. Indeed, variants that are predicted to be fitter than the wild type protein, and especially the variants *i* with 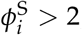, have a high *R*_*i*_ rate, which means that they circulate a lot and got fixed during viral evolution. We will deepen this point in Sections 3.6-3.7.

It is important to underline that we did not fit any parameters of our model on the SARS-CoV-2 data. Thus, this prediction as well as all the predictions presented in the following sections are truly blind predictions.

Finally, it is also instructive to analyze the localization of the variants fixed through viral evolution in the 3D structure of the spike protein. The mean values of *R*_*i*_ in the core (solvent accessibility <20%), in partially buried regions (20%-50%) and at the surface (>50%) are equal to 0.06, 0.06 and 0.23, respectively. This indicates that variants that got fixed are mainly situated in solvent-exposed regions, where they can play a key role in modulating binding with other biomolecules. Variants in buried or partially buried regions are less often observed, as these areas are more constrained from a structural point of view and are usually not involved in function.

### 3.3. Spike protein/ACE2 binding affinity and SARS-Cov-2 infectivity

We analyzed here the impact of variants on the binding of the spike protein with the ACE2 receptor. For all possible point substitutions *i* in the spike protein, we computed the change in binding affinity of the spike protein/ACE2 complex, 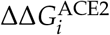, using the BeAtMuSiC program [34]. Based on the 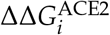values, we estimated the 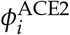viral fitness, aimed at modeling infectivity. Indeed, a higher binding affinity between the spike protein and ACE2 results in a higher efficiency of virus entry into the host’s cells [37], which in turn leads to an increase of SARS-CoV-2 infectivity.

We compared the predicted binding free energy values 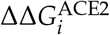with the experimentally characterized binding properties of thousands of variants introduced in the RBD of the spike protein using a yeast surface display platform, in which binding to ACE2 were quantitatively determined via flow cytometry [41]. Such deep mutagenesis scanning techniques are excellent tools to estimate biophysical quantities on a large scale. However, even though the average accuracy is reasonably good, the measured quantities are often noisy [43].

A good agreement was found between the computed 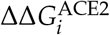 values and the large-scale measured binding affinity properties, with a Pearson’s correlation coefficient of −0.46. This result is very good, especially as not only the computed but also the experimental values have limited accuracy. It clearly underlines the quality of our prediction approach.

### 3.4. Spike protein/nAb binding affinity and immune escape

Immune evasion is the well-known mechanism used by viruses to evade from the immune system of its host, thus making its replication and spreading more efficient [44]. This mechanism involves a series of strategies such as spontaneous mutations that result in the inactivation of nAbs [45] or in the inhibition of pattern-recognition receptors initiating signalling pathways [46].

To represent the diversity of the B-cell receptor repertoire and to mimic the effect of the human immune response, we considered the set 𝒟^nAb^ of more than 30 nAbs, of which the 3D structures with the RBD of the spike protein have been experimentally resolved (see Section 2.1). We performed a large-scale *in silico* mutagenesis experiment by introducing all possible point mutations *i* in the RBD of the spike protein and by computing with BeAtMuSiC [34] the resulting change in binding free energy 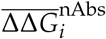 averaged over of all spike protein/nAb complexes that contain the mutation, as well as their associated fitness contribution 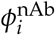 (see Eqs (2)-(4)). With this procedure, we identified key spike protein variants that are likely to either help or destroy the neutralization activity of the nAbs.

In a first stage, we performed validation tests on BeAtMuSiC’s 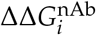 predictions. We compared them with deep mutagenesis scanning data measuring the impact of mutations in the RBD on their escape fractions from two nAbs, REGN10933 and REGN10987, which are often administrated as a cocktail to COVID-19 patients [47]. The escape fractions were estimated using a high-throughput yeast-surface-display platform, in which folded RBDs were expressed on the yeast cell surface and the fraction of cells that express mutant RBDs and that are bound to nAbs was measured [19]. Per-mutant escape fraction values close to zero indicate that the variant is bound to nAbs while values close to one indicate that it is not.

The structures of the complexes formed by the spike protein and REGN10933 or REGN10987 nAbs have recently been resolved (PDB code 6XDG). They target two different structural epitopes in the RBD of the spike protein. We did not include these structures in our set 𝒟^nAb^ as they have been resolved via cryo-EM technique at only 3.9 Å of resolution. We predicted the changes in binding affinity ΔΔ*G*_*i*_ of the two spike protein/nAb complexes caused by all RBD mutations *i* for which experimental escape fractions were available. Despite the low resolution of the 3D structures, we found very good Pearson correlation coefficients of 0.48 and 0.43 between the per-mutant escape fractions and the computed changes in affinity ΔΔ*Gi* for REGN10933 and REGN10987 nAbs, respectively.

In a second stage, we estimated the fitness contributions 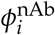 of all possible mutations *i* in the spike protein’s RBD on the basis of the predicted changes in binding free energy for the set of 31 good-resolution nAbs/spike protein complexes collected in 𝒟^nAb^. We made here and in what follows the strong approximation that these 31 nAbs represent the diversity of the human nAb repertoire. To validate this model, we compared the estimated fitness contributions 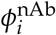 with a series of data obtained from *in vivo* experiments aimed to study the viral escape from nAbs.

We started by considering the set of 22 variants of the spike protein for which the neutralizing activity of six nAbs has been experimentally tested in terms of the relative degree of resistance (in %) of the growth of each mutant virus in the presence or in the absence of each of these nAbs [48]; we considered the average percentage over the six nAbs tested. Low percentages identify variants that escape much more from nAbs than the wild type virus and high percentages, variants that only weakly affect the wild-type spike protein/nAbs affinity. We predicted correctly 18 out of the 22 variants as having *ϕ*^nAb^ fitness values greater than one; the last four variants have *ϕ*^nAb^ 0.9. Detailed results are reported in Table 1 for the five variants shown to have the broadest *in vitro* neutralizing spectrum [48]. Our results reproduce quite well the *in vitro* trends: variants that are likely to escape from at least some nAbs tend to have fitness values larger than one. Note, moreover, that the antibodies tested in [48] are different from the nAbs of our 𝒟^nAb^ set. Because of that, we did not expect such a good match between the experiments and our predictions. This result indicates that the set 𝒟^nAb^ is truly representative of the antibody repertoire neutralizing the SARS-CoV-2 virus.

**Table 1:** List of the five variants of the spike protein RBD which have the broadest *in vitro* neutralizing spectrum, as measured in [48]. Their measured average resistance to six nAbs compared to the wild-type are given, as well as their fitness *ϕ*^nAb^ predicted on the basis of the 31 nAbs from the 𝒟^nAb^ set.

| Variants | Resistance to nAbs | $\phi^{\text{nAb}}$ |
| --- | --- | --- |
| S349A | 35 % | 1.1 |
| G446A | 37 % | 1.4 |
| G447A | 41 % | 1.6 |
| N448A | 26 % | 1.5 |
| E484A | 44 % | 1.1 |

The response to the viral infection drastically depends on the ensemble of nAbs present in the host, given that each nAb behaves differently with respect to wild-type and variant strains. In agreement with this, the predicted change in binding free energy 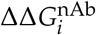 is found to strongly depend on the considered variant and nAb/spike protein complex, as clearly seen in Fig. 4. Remember that it is the average 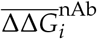 over all the nAbs that is used to define the fitness contribution *ϕ*^nAb^ and thus the overall immune escape ability.

**Figure 4.**
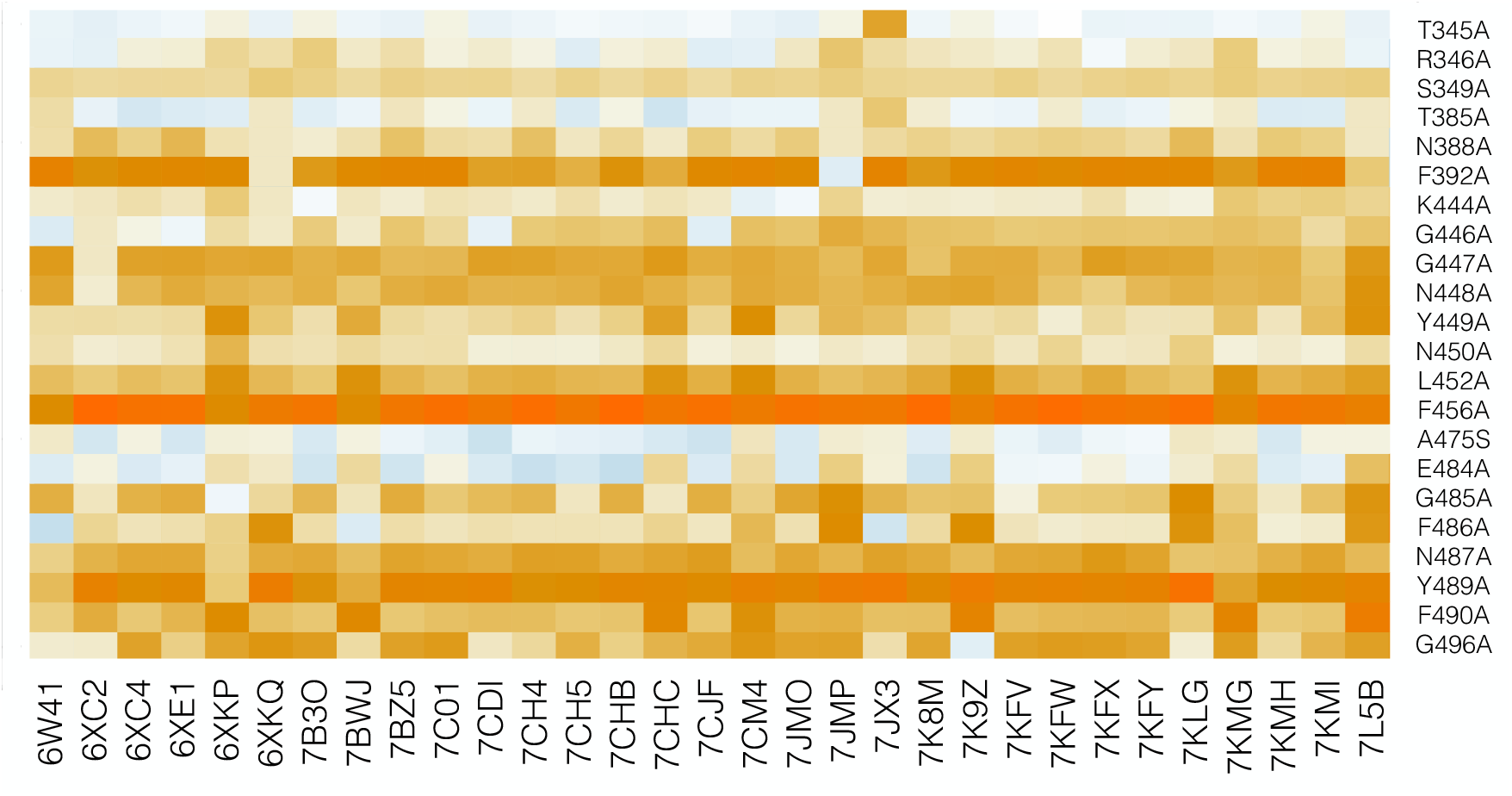
HeatMap of the predicted 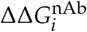values for the set of 31 nAbs/spike protein complexes from the set 𝒟^nAb^. Light blue corresponds to neutral or slightly stabilizing variants, while orange indicates mutations destabilizing the complex, which are likely to lead to the escape from the immune system.

We also validated our fitness predictions *ϕ*^nAb^ against the large-scale experimental estimation of the immune escape fractions of about 2,000 variants, averaged over a set of 17 nAbs [49]; these nAbs are not in the set 𝒟^nAb^. We found a reasonably good overall Pearson correlation coefficient of 0.29 between *ϕ*^nAb^ and measured escape fractions. Looking at more detail, the residues whose mutations most affect nAb binding belong to two regions of the RBD: the 443–450 and 484-490 loops that are situated at both sides of the ACE2 binding interface [49]. Using our set 𝒟^nAb^ of nAbs, we predicted the second region as potentially leading to immune escape with a *ϕ*^nAb^ value of 1.6. The nAb escaping capability is predicted to be weaker for the first region, with *ϕ*^nAb^ = 1.1.

A 3D representation of the per-residue fitness contributions 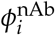in the RBD of the spike protein, averaged over all possible mutations at each position, is shown in Fig. 5. This figure is very useful to identify residues whose mutation is likely to lead to the escape from the 𝒟^nAb^ set of nAbs.

**Figure 5.**
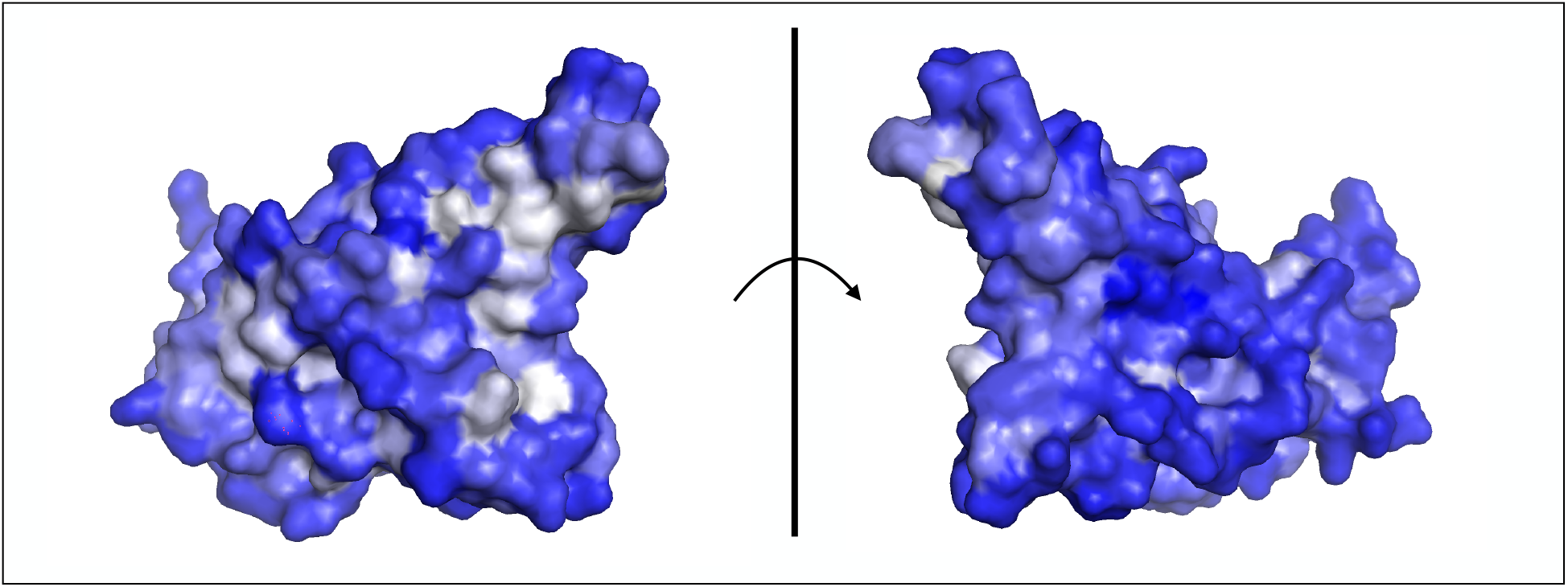
Average per-residue *ϕ*^nAb^ values mapped onto the 3D structure of the RBD of the spike protein (PDB code 6M0J). The residues whose mutations are likely to lead to the escape of SARS-CoV-2 from the nAbs are shown in white and the others in blue. The two views are related by a 180^*o*^ rotation with respect to the plane representing the ACE2 binding interface.

### 3.5. Immune escape from polyclonal human sera

We examined to what extent our method reproduces the impact of variants on the neutralizing activity of polyclonal human sera. Note that such activity depends on a wide range of factors among which inter-patient variability and time since infection [49]. Our computational approach is obviously unable to capture all intricate dependencies but rather, we expect it to detect general trends.

We used deep mutagenesis scanning data from [49], in which the escape fractions of about 2,000 single-site RBD variants were assessed on the neutralizing activity of plasma samples taken from 17 SARS-CoV-2-infected individuals, at different time points after infection. We calculated the correlation between the escape fraction for each variant averaged over the patients and post-infection time points and the predicted fitness contributions *ϕ*^nAb^ computed from the 𝒟^nAb^ set of nAbs. We obtained a reasonably good Pearson correlation coefficient of 0.35 between the predicted and measured quantities.

Only few residues appear to contribute substantially to the escape mechanisms, when averaged over the whole plasma sample collection. Indeed, only 23 residues have an average escape fraction greater than 3%. Our predictions for these residues are in very good agreement with experiments: we obtained an average per-residue *ϕ*^nAb^ equal to 1.5. Residue F456 shows almost perfect agreement: it has the highest measured escape fraction, and also has the highest predicted *ϕ*^nAb^ value, equal to 2.2. Almost all substitutions at that position are predicted to strongly impact on the binding in the majority of spike-protein/nAbs complexes analyzed.

Finally, it is interesting to compare the measured immune escaping fractions in polyclonal plasma discussed in this section with the experimentally characterized escape fractions in the set of nAbs studied in [49] and discussed in Section 3.4. We found that their linear correlation coefficient is equal to 0.4, which indicates there are differences between the tested cocktail of nAbs and serum plasma. Possible explanations include the scarcity of the tested antibodies in the polyclonal plasma, or the subdominance of the epitopes they target [49].

### 3.6. Overall variant fitness, transmissibility, infectivity and immune escape

We focused on five SARS-CoV-2 variants most frequently observed worldwide, as reported in the GISAID database [42] in March 2021, and predicted their fitness; the results are shown in Table 2.

**Table 2:**
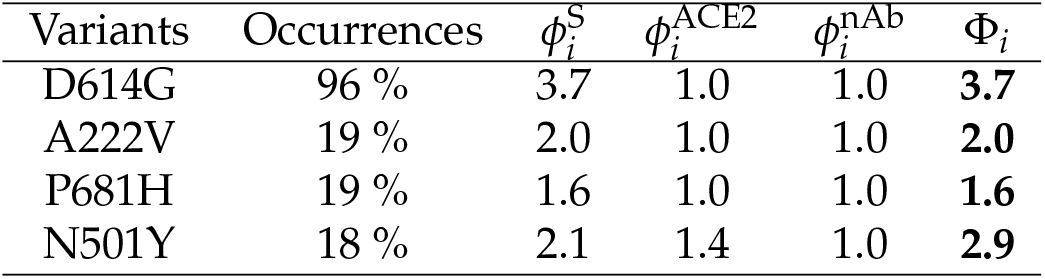
The five most widely observed variants and their predicted fitness. Occurrences refer to their number of occurrences in the GISAID database [42], 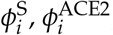, and 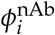 to the fitness contributions of the variants *i* related to the stability of the spike protein, its binding affinity for ACE2 and its escape propensity from the host’s immune system, respectively, and Φ_*i*_ to the total fitness.

The most frequently observed spike protein variant involves the substitution of aspartic acid at position 614 into glycine, situated outside the RBD. This variant quickly became dominant after its appearance in early 2020 [36,50]. We correctly predicted a substantial increase of fitness for this variant with respect to wild type, which is driven by an increased stability of the spike protein 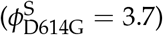. We hypothesize that this stabilization leads to a higher person-to-person viral transmissibility, as also suggested in [36,50,51] and observed *in vivo* [51]. In the latter study, a stabilization of the spike protein was measured upon D614G substitution via a strengthening of the S1-S2 subunit interactions, where S1 is the receptor binding subunit containing the RBD and S2 is the membrane fusion subunit. In contrast, this variant was shown to alter neither the binding of the spike protein to ACE2 nor the antibody neutralization, as it is situated outside the RBD [51]. We also correctly reproduced this result, with fitness values of 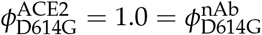 (Table 2). The overall predicted fitness is thus Φ_D614G_ = 3.7.

Two other variants, A222V and P681H, show similar albeit less pronounced trends. Our results predict an increase in transmissibility 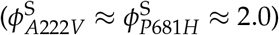, but to a lesser extent than D614G. Experimental data are in agreement with the weaker impacts of these variants on the spike protein fitness and the viral transmissibility compared to D614G [52,53]. The A222V variant has been related to the large outbreaks in Europe in early summer 2020, while P681H is associated to the so-called UK lineage (B.1.1.7) that appeared in UK in late 2020 and is now becoming dominant in Europe in the current outbreaks.

Finally, N501Y is also a widely spread variant appearing in all major lineages, *i*.*e*. UK (B.1.1.7), Brazilian (P.1) and South African (B.1.351) lineages. We predicted this variant as having a high overall fitness Φ due to a combination of increased fitness contributions 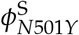 and 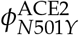, but a 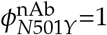. In other words, we predicted this variant to be more transmissible and infectious than the wild type but to have no impact on the response of the human immune system. More precisely, we predicted N501Y as improving the stability of the spike protein RBD and its binding affinity for ACE2; the latter property is also suggested by another computational study [54]. No clinical data suggest that N501Y is able to escape from the immune post-vaccination response [55], which tends to support our prediction results.

### 3.7. Viral evolution and overall fitness

We applied our prediction pipeline to analyze SARS-CoV-2 evolution, focusing on the spike protein. We started by predicting the viral fitness F of all the SARS-CoV-2 strains collected in the GISAID database from December 2019 till March 2021, which amounts to about 7.8 × 10^5^ strains. We subdivided the strains according to the month of collection and computed the per-month average of viral fitness. The results are reported in Fig. 6.a as a function of time. Clearly, we predict an increase of the viral fitness since the beginning of the infection in December 2019, in agreement with epidemiological results. This result once again demonstrates the quality of our computational pipeline.

**Figure 6.**
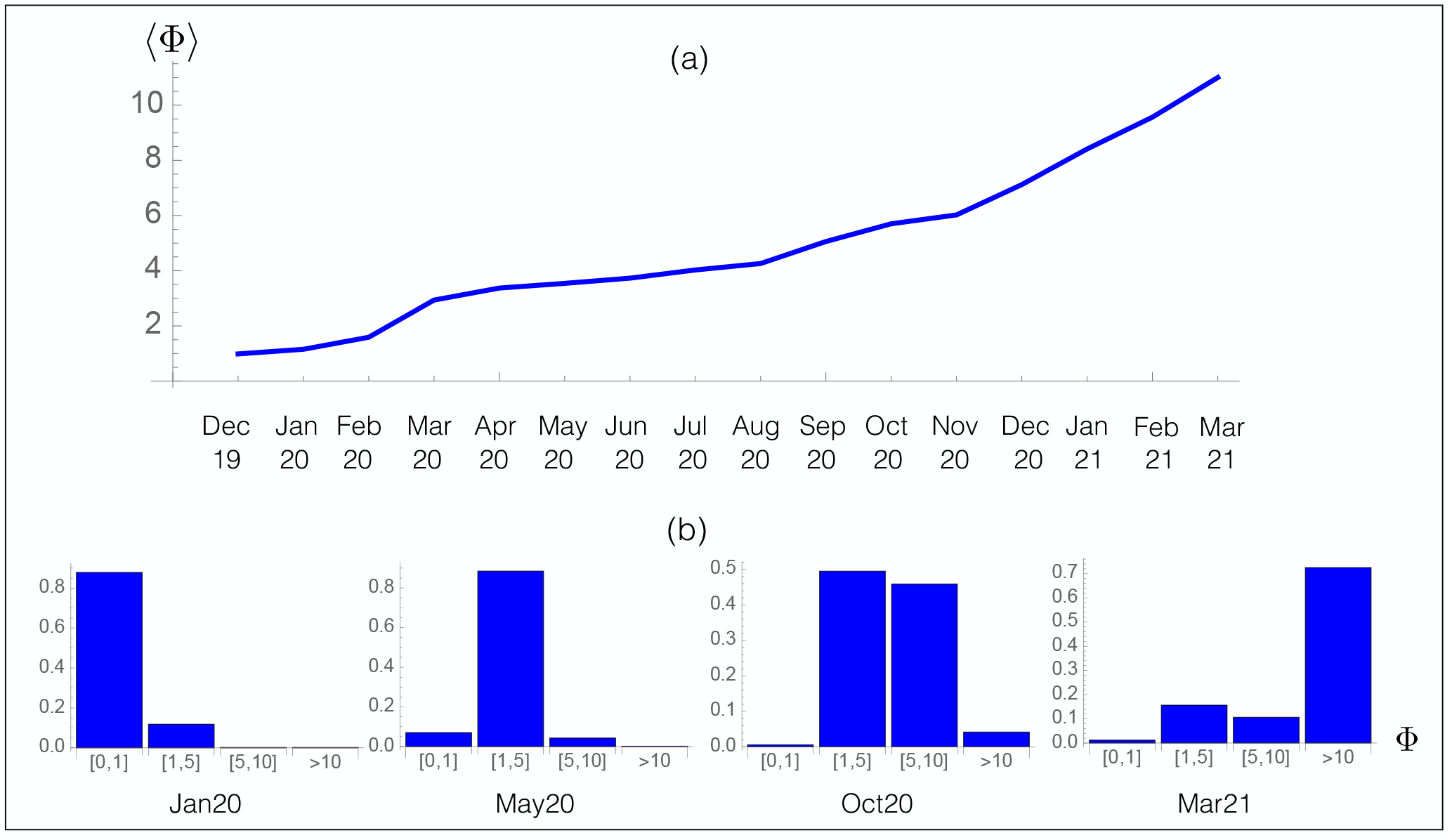
Time evolution of the predicted overall fitness Φ. (a) Average fitness ⟨Φ ⟩ per month for the SARS-CoV-2 strains collected from the GISAID database as a function of time; (b) Probability distribution of Φ for the SARS-CoV-2 strains collected from GISAID during a given month.

Note that to predict the future evolution of the fitness Φ, it is necessary to take into account different parameters such as the varying repertoire of human nAbs and the effect of vaccination. While the fitness contributions *ϕ*^S^ and *ϕ*^ACE2^ are expected to reach a plateau when the spike protein sequence becomes optimal for stability and for binding to ACE2, the cat-and-mouse game played by the virus and its host leads the host to continuously adapt its B-cell repertoire to the new variants of the virus, so that *ϕ*^nAb^ certainly increases with respect to the old nAbs, but not with respect to the new nAbs. In total, the overall fitness Φ is expected to plateau after some time, or at least increase less.

We analyzed in more detail the evolution of the partial distribution function of the per-month averaged fitness in Fig. 6.b. In January 2020, the population was dominated by the wild type strain whose fitness F is by definition equal to one. The effect of the D614G spike protein variant with a predicted Φ ∼ 4 is observed from May 2020, while in October of the same year, aΔitional mutations with Φ = 2.0, such as A222V, started to be fixed in the population, leading to a further increase of F. In March 2021, the distribution became dominated by new variants, *i*.*e*. UK, South-African and Brazilian variants, with a much higher fitness than both the wild-type and D614G strains.

Finally, we carefully checked that our large-scale mutagenesis predictions are not biased towards high fitness values. Indeed, such bias could potentially cause a trivial increase in fitness upon evolution and lead to erroneous interpretations. To verify this, we created 2 × 10^6^ viral strains by inserting either three or five random mutations in the wild-type spike protein and assumed that they got fixed with probability one independently of their fitness value; the number of random mutations was chosen based on the average number of single variants per strain in the GISAID database which is between three and four. We then computed the fitness Φ for all these random variant strains. On the other hand, we plotted the Φ distribution of the real variant strains observed in the GISAID database. The fitness distribution of the two simulated and the real viral strains are completely different, as shown in Fig. 7. When three or five random mutations are inserted in the spike protein, the Φ distributions have a median value of 0.32 and 0.12, respectively; moreover, 79% and 86% of the mutated strains have a lower overall fitness Φ than the wild-type virus. In contrast, the distribution of real strains has a median of 4.8 and basically all the strains (99.5%) have a predicted fitness higher than the wild type. This analysis further supports the unbiased nature and validity of our computational approach.

**Figure 7.**
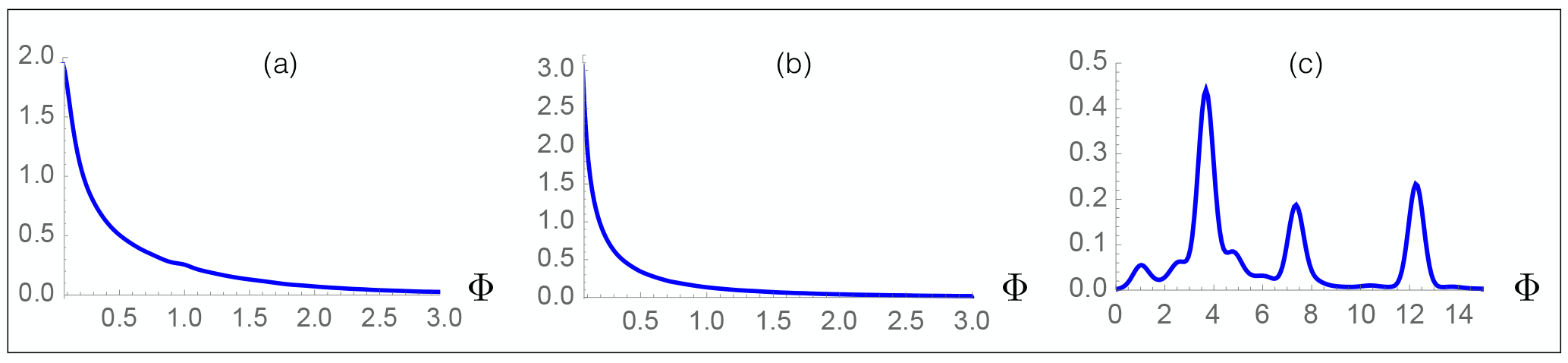
Probability distribution of the predicted fitness Φ for 10^6^ simulated viral strains obtained by inserting (a) three and (b) five random mutations in the spike protein; (c) Probability distribution function of the predicted fitness Φ for all the strains collected in the GISAID database.

## 4. Conclusion

Here we set up and validated SpikePro, a simple computational model that predicts the impact of spike protein variants on the SARS-CoV-2 fitness and more specifically, on viral transmissibility, infectivity and ability of escaping from the host’s immune system. Moreover, the program is easy to use and can be freely downloaded from github.com/3BioCompBio/SpikeProSARS-CoV-2. SpikePro allows identifying, with good accuracy and in a few seconds, new SARS-CoV-2 variants with high fitness which need to be closely monitored by health agencies. It has thus a central role to play in the genomic surveillance programs of the new SARS-CoV-2 strains, especially in the coming future with the growing number of people vaccinated and thus the larger selective pressure on the virus [56].

We thoroughly analyzed and validated SpikePro on a wide series of experimental, epidemiological and clinical data available. Despite the simplicity of the model, the approximations made, and the absence of parameters that were fitted to optimize the accuracy of the predictions, the SpikePro pipeline reproduces well the collected data. Whether the validation is performed on large-scale mutagenesis data, nAb cocktails or polyclonal human sera, whether the comparison involves the fitness of the spike protein, of the spike protein/ACE2 complex, or of a series of spike protein/nAb complexes, the results are very good with correlation coefficients in the 0.3 to 0.5 range.

In addition, SpikePro predicts a high overall fitness value for the frequently occurring variants such as the UK, Brazilian or South-African variants and correctly identifies the main fitness contributions. It also reproduces quite well the overall fitness evolution of the SARS-CoV-2 virus over the past pandemic year.

It has to be emphasized that the SpikePro model, besides being able to reproduce known results, has a true prediction potential in describing and interpreting the effect of new spike protein variants that could be fixed in the near future and the future SARS-CoV-2 evolution, owing to the physical description of the fitness in terms of free energy contributions, which are estimated using the well-known structure-based PoPMuSiC and BeAtMuSiC predictors [33,34].

Despite the progress we made towards a better understanding of the molecular mechanisms underlying the SARS-CoV-2 fitness, we made some approximations in the construction of our model which we will try to relax in future studies. For example, we did not take into account possible amino acid deletions or insertions in the spike protein, although they certainly influence the viral fitness. It would also be interesting to take into account epistatic effects. Indeed, while more and more variants get fixed, interactions between them are expected to become non-negligible. Furthermore, the model should be extended to other proteins of the SARS-CoV-2 virus such as the non-structural protein 1 (Nsp1) which also contributes to immune evasion [57], rather than considering the spike protein only. Finally, when more nAbs/spike protein complexes will be resolved at high resolution, they will enrich our set 𝒟^nAb^ and better describe the B-cell receptor repertoire. Considering a weighted combination of the effects of RBD variants on all nAbs depending on different factors such as time and vaccination status would further improve our method in mimicking the immune response and its temporal evolution.

## Author Contributions

Conceptualization, F.P. and M.R.; formal analysis and investigation F.P. and M.R.; methodology and validation, F.P.; writing–original draft preparation, F.P. and M.R.; writing–review and editing F.P. and M.R. All authors have read and agreed to the published version of the manuscript.

## Acknowledgements

FP and MR are Postdoctoral Researcher and Research Director, respectively, at the F.R.S.-FNRS Fund for Scientific Research.

## Data availability

The SpikePro algorithm is freely available on GitHub (https://github.com/3BioCompBio/SpikeProSARS-CoV-2).

## References

1. Dong E.; Du, H.; Gardner, L. An interactive web-based dashboard to track COVID-19 in real time. The Lancet infectious diseases 2020, 20, 533–534.

2. Sanders, J.M.; Monogue, M.L.; Jodlowski, T.Z.; Cutrell, J.B. Pharmacologic treatments for coronavirus disease 2019 (COVID-19): a review. Jama 2020, 323, 1824–1836.

3. Le, T.T.; Andreadakis, Z.; Kumar, A.; Román, R.G.; Tollefsen, S.; Saville, M.; Mayhew, S.; others. The COVID-19 vaccine development landscape. Nat Rev Drug Discov 2020, 19, 305–306.

4. Baden, L.R.; El Sahly, H.M.; Essink, B.; Kotloff, K.; Frey, S.; Novak, R.; Diemert, D.; Spector, S.A.; Rouphael, N.; Creech, C.B.; others. Efficacy and safety of the mRNA-1273 SARS-CoV-2 vaccine. New England Journal of Medicine 2020.

5. Polack, F.P.; Thomas, S.J.; Kitchin, N.; Absalon, J.; Gurtman, A.; Lockhart, S.;Perez, J.L.; Pérez Marc, G.; Moreira, E.D.; Zerbini, C.; others. Safety and efficacy of the BNT162b2 mRNA Covid-19 vaccine. New England Journal of Medicine 2020, 383, 2603–2615.

6. Voysey, M.; Clemens, S.A.C.; Madhi, S.A.; Weckx, L.Y.; Folegatti, P.M.; Aley, P.K.; Angus, B.; Baillie, V.L.; Barnabas, S.L.; Bhorat, Q.E.; others. Safety and efficacy of the ChAdOx1 nCoV-19 vaccine (AZD1222) against SARS-CoV-2: an interim analysis of four randomised controlled trials in Brazil, South Africa, and the UK. The Lancet 2021, 397, 99–111.

7. Logunov, D.Y.; Dolzhikova, I.V.; Shcheblyakov, D.V.; Tukhvatulin, A.I.; Zubkova, O.V.; Dzharullaeva, A.S.; Kovyrshina, A.V.; Lubenets, N.L.; Grousova, D.M.; Erokhova, A.S.; others. Safety and efficacy of an rAd26 and rAd5 vector-based heterologous prime-boost COVID-19 vaccine: an interim analysis of a randomised controlled phase 3 trial in Russia. The Lancet 2021.

8. Sadoff, J.; Le Gars, M.; Shukarev, G.; Heerwegh, D.; Truyers, C.; de Groot, A.M.; Stoop, J.; Tete, S.; Van Damme, W.; Leroux-Roels, I.; others. Interim Results of a Phase 1–2a Trial of Ad26. COV2. S Covid-19 Vaccine. New England Journal of Medicine 2021.

9. Keech, C.; Albert, G.; Cho, I.; Robertson, A.; Reed, P.; Neal, S.; Plested, J.S.; Zhu, M.; Cloney-Clark, S.; Zhou, H.; others. Phase 1–2 trial of a SARS-CoV-2 recombinant spike protein nanoparticle vaccine. New England Journal of Medicine 2020, 383, 2320–2332.

10. Bhimraj, A.; Morgan, R.L.; Shumaker, A.H.; Lavergne, V.; Baden, L.; Cheng, V.C.C.; Edwards, K.M.; Gandhi, R.; Muller, W.J.; O’Horo, J.C.; others. Infectious Diseases Society of America guidelines on the treatment and management of patients with COVID-19. Clinical Infectious Diseases 2020.

11. Weinreich, D.M.; Sivapalasingam, S.; Norton, T.; Ali, S.; Gao, H.; Bhore, R.; Musser, B.J.; Soo, Y.; Rofail, D.; Im, J.; others. REGN-COV2, a Neutralizing Antibody Cocktail, in Outpatients with Covid-19. New England Journal of Medicine 2020.

12. Jiang, S.; Zhang, X.; Yang, Y.; Hotez, P.J.; Du, L. Neutralizing antibodies for the treatment of COVID-19. Nature Biomedical Engineering 2020, 4, 1134–1139.

13. Joyner, M.J.; Carter, R.E.; Senefeld, J.W.; Klassen, S.A.; Mills, J.R.; Johnson, P.W.; Theel, E.S.; Wiggins, C.C.; Bruno, K.A.; Klompas, A.M.; others. Convalescent plasma antibody levels and the risk of death from covid-19. New England Journal of Medicine 2021.

14. Alcami, A.; Koszinowski, U.H. Viral mechanisms of immune evasion. Trends in microbiology 2000, 8, 410–418.

15. Williams, T.C.; Burgers, W.A. SARS-CoV-2 evolution and vaccines: cause for concern? The Lancet Respiratory Medicine 2021.

16. Ku, Z.; Xie, X.; Davidson, E.; Ye, X.; Su, H.; Menachery, V.D.; Li, Y.; Yuan, Z.; Zhang, X.; Muruato, A.E.; others. Molecular determinants and mechanism for antibody cocktail preventing SARS-CoV-2 escape. Nature communications 2021, 12, 1–13.

17. Weisblum, Y.; Schmidt, F.; Zhang, F.; DaSilva, J.; Poston, D.; Lorenzi, J.C.; Muecksch, F.; Rutkowska, M.; Hoffmann, H.H.; Michailidis, E.; others. Escape from neutralizing antibodies by SARS-CoV-2 spike protein variants. Elife 2020, 9, e61312.

18. Greaney, A.J.; Starr, T.N.; Gilchuk, P.; Zost, S.J.; Binshtein, E.; Loes, A.N.; Hilton, S.K.; HuΔleston, J.; Eguia, R.; Crawford, K.H.; others. Complete mapping of mutations to the SARS-CoV-2 spike receptor-binding domain that escape antibody recognition. Cell host & microbe 2021, 29, 44–57.

19. Starr, T.N.; Greaney, A.J.; AΔetia, A.; Hannon, W.W.; Choudhary, M.C.; Dingens, A.S.; Li, J.Z.; Bloom, J.D. Prospective mapping of viral mutations that escape antibodies used to treat COVID-19. Science 2021.

20. McCarthy, K.R.; Rennick, L.J.; Nambulli, S.; Robinson-McCarthy, L.R.; Bain, W.G.; Haidar, G.; Duprex, W.P. Recurrent deletions in the SARS-CoV-2 spike glycoprotein drive antibody escape. Science 2021. doi:10.1126/science.abf6950.

21. Tegally, H.; Wilkinson, E.; Giovanetti, M.; Iranzadeh, A.; Fonseca, V.; Giandhari, J.; Doolabh, D.; Pillay, S.; San, E.J.; Msomi, N.; others. Emergence and rapid spread of a new severe acute respiratory syndrome-related coronavirus 2 (SARS-CoV-2) lineage with multiple spike mutations in South Africa. medRxiv 2020.

22. Andreano, E.; Piccini, G.; Licastro, D.; Casalino, L.; Johnson, N.V.; Paciello, I.; Dal Monego, S.; Pantano, E.; Manganaro, N.; Manenti, A.; others. SARS-CoV-2 escape in vitro from a highly neutralizing COVID-19 convalescent plasma. bioRxiv 2020.

23. Wibmer, C.K.; Ayres, F.; Hermanus, T.; Madzivhandila, M.; Kgagudi, P.; Lambson, B.E.; Vermeulen, M.; van den Berg, K.; Rossouw, T.; Boswell, M.; others. SARS-CoV-2 501Y. V2 escapes neutralization by South African COVID-19 donor plasma. BioRxiv.

24. Wang, Z.; Schmidt, F.; Weisblum, Y.; Muecksch, F.; Barnes, C.O.; Finkin, S.; Schaefer-Babajew, D.; Cipolla, M.; Gaebler, C.; Lieberman, J.A.; others. mRNA vaccine-elicited antibodies to SARS-CoV-2 and circulating variants. bioRxiv 2021.

25. Collier, D.A.; De Marco, A.; Ferreira, I.A.; Meng, B.; Datir, R.; Walls, A.C.; Kemp S, S.A.; Bassi, J.; Pinto, D.; Fregni, C.S.; Bianchi, S.; Tortorici, M.A.; Bowen, J.; Culap, K.; Jaconi, S.; Cameroni, E.; Snell, G.; Pizzuto, M.S.; Pellanda, A.F.; Garzoni, C.; Riva, A.; .; Elmer, A.; Kingston, N.; Graves, B.; McCoy, L.E.; Smith, K.G.; Bradley, J.R.; James Thaventhiran, J.; Lourdes Ceron-Gutierrez, L.; Barcenas-Morales, G.; Virgin, H.W.; Lanzavecchia, A.; Piccoli, L.; Doffinger, R.; Wills, M.; Veesler, D.; Corti, D.; Gupta, R.K. SARS-CoV-2 B.1.1.7 escape from mRNA vaccine-elicited neutralizing antibodies. medRxiv 2021. doi:10.1101/2021.01.19.21249840.

26. Thomson, E.C.; Rosen, L.E.; Shepherd, J.G.; Spreafico, R.; da Silva Filipe, A.; Wojcechowskyj, J.A.; Davis, C.; Piccoli, L.; Pascall, D.J.; Dillen, J.; others. Circulating SARS-CoV-2 spike N439K variants maintain fitness while evading antibody-mediated immunity. Cell 2021.

27. Lauring, A.S.; Hodcroft, E.B. Genetic Variants of SARS-CoV-2—What Do They Mean? JAMA.

28. Berman, H.; Westbrook, J.; Feng, Z.; Gilliland, G.; Bhat T.; H., W.; Shindyalov, I.; Bourne, P. The Protein Data Bank. Nucleic Acids Research 2000, 28, 235–242.

29. Walls, A.C.; Park, Y.J.; Tortorici, M.A.; Wall, A.; McGuire, A.T.; Veesler, D. Structure, function, and antigenicity of the SARS-CoV-2 spike glycoprotein. Cell 2020, 181, 281–292.

30. Waterhouse, A.; Bertoni, M.; Bienert, S.; Studer, G.; Tauriello, G.; Gumienny, R.; Heer, F.T.; de Beer, T.A.P.; Rempfer, C.; Bordoli, L.; others. SWISS-MODEL: homology modelling of protein structures and complexes. Nucleic acids research 2018, 46, W296–W303.

31. Lan, J.; Ge, J.; Yu, J.; Shan, S.; Zhou, H.; Fan, S.; Zhang, Q.; Shi, X.; Wang, Q.; Zhang, L.; others. Structure of the SARS-CoV-2 spike receptor-binding domain bound to the ACE2 receptor. Nature 2020, 581, 215–220.

32. Raybould, M.I.; Kovaltsuk, A.; Marks, C.; Deane, C.M. CoV-AbDab: the coronavirus antibody database. BioRxiv 2020.

33. Dehouck, Y.; Grosfils, A.; Folch, B.; Gilis, D.; Bogaerts, P.; Rooman, M. Fast and accurate predictions of protein stability changes upon mutations using statistical potentials and neural networks: PoPMuSiC-2.0. Bioinformatics 2009, 25, 2537–2543.

34. Dehouck, Y.; Kwasigroch, J.M.; Rooman, M.; Gilis, D. BeAtMuSiC: prediction of changes in protein–protein binding affinity on mutations. Nucleic acids research 2013, 41, W333–W339.

35. Wargo, A.R.; Kurath, G. Viral fitness: definitions, measurement, and current insights. Current opinion in virology 2012, 2, 538–545.

36. Baric, R.S. Emergence of a Highly Fit SARS-CoV-2 Variant. New England Journal of Medicine 2020.

37. Shang, J.; Ye, G.; Shi, K.; Wan, Y.; Luo, C.; Aihara, H.; Geng, Q.; Auerbach, A.; Li, F. Structural basis of receptor recognition by SARS-CoV-2. Nature 2020, 581, 221–224.

38. Echave, J.; Jackson, E.L.; Wilke, C.O. Relationship between protein thermodynamic constraints and variation of evolutionary rates among sites. Physical biology 2015, 12, 025002.

39. Pucci, F.; Bernaerts, K.; Teheux, F.; Gilis, D.; Rooman, M. Symmetry principles in optimization problems : an application to protein stability prediction. IFAC-PapersOnLine 2015, 48, 458—-463.

40. Pucci, F.; Bernaerts, K.V.; Kwasigroch, J.M.; Rooman, M. Quantification of biases in predictions of protein stability changes upon mutations. Bioinformatics 2018, 34, 3659–3665.

41. Starr, T.N.; Greaney, A.J.; Hilton, S.K.; Ellis, D.; Crawford, K.H.; Dingens, A.S.; Navarro, M.J.; Bowen, J.E.; Tortorici, M.A.; Walls, A.C.; others. Deep mutational scanning of SARS-CoV-2 receptor binding domain reveals constraints on folding and ACE2 binding. Cell 2020, 182, 1295–1310.

42. Elbe, S.; Buckland-Merrett, G. Data, disease and diplomacy: GISAID’s innovative contribution to global health. Global Challenges 2017, 1, 33–46.

43. Fowler, D.M.; Stephany, J.J.; Fields, S. Measuring the activity of protein variants on a large scale using deep mutational scanning. Nature protocols 2014, 9, 2267–2284.

44. Beachboard, D.C.; Horner, S.M. Innate immune evasion strategies of DNA and RNA viruses. Current opinion in microbiology 2016, 32, 113–119.

45. Doud, M.B.; Hensley, S.E.; Bloom, J.D. Complete mapping of viral escape from neutralizing antibodies. PLoS pathogens 2017, 13, e1006271.

46. Bowie, A.G.; Unterholzner, L. Viral evasion and subversion of pattern-recognition receptor signalling. Nature Reviews Immunology 2008, 8, 911–922.

47. Hansen, J.; Baum, A.; Pascal, K.E.; Russo, V.; Giordano, S.; Wloga, E.; Fulton, B.O.; Yan, Y.; Koon, K.; Patel, K.; others. Studies in humanized mice and convalescent humans yield a SARS-CoV-2 antibody cocktail. Science 2020, 369, 1010–1014.

48. Ku, Z.; Xie, X.; Davidson, E.; Ye, X.; Su, H.; Menachery, V.D.; Li, Y.; Yuan, Z.; Zhang, X.; Muruato, A.E.; i Escuer, A.G.; Tyrell, B.; Doolan, K.; Doranz, B.J.; Wrapp, D.; Bates, P.F.; McLellan, J.S.; Weiss, S.R.; Zhang, N.; Shi, P.Y.; An, Z. Molecular determinants and mechanism for antibody cocktail preventing SARS-CoV-2 escape. Nature Communications 2021, 12, 469.

49. Greaney, A.J.; Loes, A.N.; Crawford, K.H.; Starr, T.N.; Malone, K.D.; Chu, H.Y.; Bloom, J.D. Comprehensive mapping of mutations in the SARS-CoV-2 receptor-binding domain that affect recognition by polyclonal human plasma antibodies. Cell host & microbe 2021.

50. Hou, Y.J.; Chiba, S.; Halfmann, P.; Ehre, C.; Kuroda, M.; Dinnon, K.H.; Leist, S.R.; Schäfer, A.; Nakajima, N.; Takahashi, K.; Lee, R.E.; Mascenik, T.M.; Graham, R.; Edwards, C.E.; Tse, L.V.; Okuda, K.; Markmann, A.J.; Bartelt, L.; de Silva, A.; Margolis, D.M.; Boucher, R.C.; Randell, S.H.; Suzuki, T.; Gralinski, L.E.; Kawaoka, Y.; Baric, R.S. SARS-CoV-2 D614G variant exhibits efficient replication ex vivo and transmission in vivo. Science 2020, 370, 1464–1468. doi:10.1126/science.abe8499.

51. Zhang, L.; Jackson, C.B.; Mou, H.; Ojha, A.; Peng, H.; Quinlan, B.D.; Rangarajan, E.S.; Pan, A.; Vanderheiden, A.; Suthar, M.S.; Li, W.; Izard, T.; Rader, C.; Farzan, M.; Choe, H. SARS-CoV-2 spike-protein D614G mutation increases virion spike density and infectivity. Nature Communications 2020, 11, 6013.

52. Hodcroft, E.B.; Zuber, M.; Nadeau, S.; Crawford, K.H.D.; Bloom, J.D.; Veesler, D.; Vaughan, T.G.; Comas, I.; Candelas, F.G.; .; Stadler, T.; Neher, R.A. Emergence and spread of a SARS-CoV-2 variant through Europe in the summer of 2020. medRxiv 2020. doi:10.1101/2020.10.25.20219063.

53. Peacock, T.P.; Goldhill, D.H.; Zhou, J.; Baillon, L.; Frise, R.; Swann, O.C.; Kugathasan, R.; Penn, R.; Brown, J.C.; Sanchez-David, R.Y.; Braga, L.; Williamson, M.K.; Hassard, J.A.; Staller, E.; Hanley, B.; Osborn, M.; Giacca, M.; Davidson, A.D.; Matthews, D.A.; Barclay, W.S. The furin cleavage site of SARS-CoV-2 spike protein is a key determinant for transmission due to enhanced replication in airway cells. bioRxiv 2020. doi:10.1101/2020.09.30.318311.

54. Villoutreix, B.O.; Calvez, V.; Marcelin, A.G.; Khatib, A.M. In Silico Investigation of the New UK (B.1.1.7) and South African (501Y.V2) SARS-CoV-2 Variants with a Focus at the ACE2–Spike RBD Interface. International Journal of Molecular Sciences 2021, 22. doi:10.3390/ijms22041695.

55. Xie, X.; Liu, Y.; Liu, J.; Zhang, X.; Zou, J.; Fontes-Garfias, C.R.; Xia, H.; Swanson, K.A.; Cutler, M.; Cooper, D.; Menachery, V.D.; Weaver, S.C.; Dormitzer, P.R.; Shi, P.Y. Neutralization of SARS-CoV-2 spike 69/70 deletion, E484K and N501Y variants by BNT162b2 vaccine-elicited sera. Nature Medicine 2021.

56. Burioni, R.; Topol, E.J. Assessing the human immune response to SARS-CoV-2 variants. Nature Medicine 2021, pp. 1–2.

57. Thoms, M.; Buschauer, R.; Ameismeier, M.; Koepke, L.; Denk, T.; Hirschenberger, M.; Kratzat, H.; Hayn, M.; Mackens-Kiani, T.; Cheng, J.; others. Structural basis for translational shutdown and immune evasion by the Nsp1 protein of SARS-CoV-2. Science 2020, 369, 1249–1255.

